# Whole genomic sequencing as a tool for diagnosis of drug and multidrug-resistance tuberculosis in an endemic region in Mexico

**DOI:** 10.1101/551481

**Authors:** Carlos Francisco Madrazo-Moya, Irving Cancino-Muñoz, Betzaida Cuevas-Cordoba, Vanessa Gonzalez-Covarrubias, Martín Barbosa-Amezcua, Xavier Soberón, Raquel Muñiz-Salazar, Armando Martínez-Guarneros, Claudia Backer, José Zarrabal-Meza, Clara Sampieri-Ramirez, Antonio Enciso-Moreno, Michael Lauzardo, Iñaki Comas, Roberto Zenteno-Cuevas

**Author notes:** These authors contributed to conceptualization, data analysis and writing, review and editing of manuscript (Carlos Francisco Madrazo-Moya, Irving Cancino-Muñoz, Raquel Muñiz-Salazar, Clara Sampieri-Ramirez, Antonio Enciso-Moreno, Iñaki Comas, Roberto Zenteno-Cuevas). These authors contributed to the provision of study materials and methodology (Irving Cancino-Muñoz, Betzaida Cuevas-Cordoba, Vanessa Gonzalez-Covarrubias, Martín Barbosa-Amezcua, Xavier Soberón, Armando Martínez-Guarneros, Claudia Backer, José Zarrabal-Meza, Michael Lauzardo, Iñaki Comas). Corresponding author: (RZ-C).

## Abstract

**Background:** Whole genome sequencing (WGS) has been proposed as a tool for diagnosing drug resistance in tuberculosis. However, reports of its effectiveness in endemic countries with important numbers of drug resistance are scarce. The goal of this study was to evaluate the effectiveness of this procedure in isolates from a tuberculosis endemic region in Mexico.

**Methods:** WGS analysis was performed in 81 tuberculosis positive clinical isolates with a known phenotypic profile of resistance against first-line drugs (isoniazid, rifampin, ethambutol, pyrazinamide and streptomycin). Mutations related to drug resistance were identified for each isolate; drug resistant genotypes were predicted and compared with the phenotypic profile. Genotypes and transmission clusters based on genetic distances were also characterized.

**Findings:** Prediction by WGS analysis of resistance against isoniazid, rifampicin, ethambutol, pyrazinamide and streptomycin showed sensitivity values of 84%, 96%, 71%, 75% and 29%, while specificity values were 100%, 94%, 90%, 90% and 98%, respectively. Prediction of multidrug resistance showed a sensitivity of 89% and specificity of 97%. Moreover, WGS analysis revealed polymorphisms related to second-line drug resistance, enabling classification of eight and two clinical isolates as pre- and extreme drug-resistant cases, respectively.

Four lineages were identified in the population (L1, L2, L3 and L4). The most frequent of these was L4, which included 90% (77) of the isolates. Six transmission clusters were identified; the most frequent was TC6, which included 13 isolates with a L4.1.1 and a predominantly multidrug-resistant condition.

**Conclusion:** The results illustrate the utility of WGS for establishing the potential for prediction of resistance against first and second line drugs in isolates of tuberculosis from the region. They also demonstrate the feasibility of this procedure for use as a tool to support the epidemiological surveillance of drug- and multidrug-resistant tuberculosis.

## Introduction

According to the annual report of the World Health Organization in 2017, 10.5 million new cases and 1.7 million deaths were related to tuberculosis (TB), establishing TB as the infectious disease with the greatest impact on human health. [1]

Close to 25% of TB cases recorded worldwide every year show resistance (DR-TB) to at least one of the four antibiotics used in first line treatment against this disease; Rifampicin (R), Isoniazid (H), Ethambutol (E) and Pyrazinamide (Z). Of these DR-TB isolates, about 5% evolve into multi-drug resistant tuberculosis; i.e., they present a combined resistance to isoniazid and rifampicin (MDR-TB), which may evolve into an aggravated form known as extreme drug-resistant tuberculosis (TB-XDR), which are isolates that possess simultaneous resistance to a flouroquinolone and at least one of the three second-line injectable drugs (amikacin, kanamycin and capreomicin). The number of isolates with these characteristics is steadily increasing, reducing the possibility of eliminating TB as a global public health problem by 2020. [2]

Whole Genome Sequencing (WGS) provides accurate information about polymorphisms, insertions and deletions (indels) of potential relevance to the rapid prediction of drug susceptibility phenotypes of clinical importance [3,4]. The implementation of WGS has potential for improving traditional phenotypic procedures to detect DR-TB cases and to influence clinical decision-making in high-income settings [5–11], in addition, this technique has also been successfully used to examine transmission dynamics, identify transmission clusters and reconstruct timelines of acquisition of antimicrobial resistance [12–15].

The current body of knowledge regarding mutations that confer drug resistance has led to the suggestion that identification of true DR-TB isolates may be more complex than previously thought [16], raising the need to further this type of study in order to identify the changes involved and increase diagnostic efficiency. Moreover, there is a lack of evidence regarding the utility of WGS in emerging or poorly-developed countries, which are precisely those most affected by drug-resistant tuberculosis, and where information relating to the polymorphism associated with drug resistance is limited [6,17–21]

In Mexico, tuberculosis is endemic. According to the 2017 Global Report, 22,869 new cases of TB were reported in 2016, with an incidence of 22 cases per 100,000 inhabitants [1]. DR-TB cases were observed in 2.5% of these cases, and there was also a prevalent population of 610 individuals with MDR-TB. Moreover, increasing numbers of XDR-TB cases are reported every year [22]. These figures make Mexico the country in the Americas with the third highest contribution of tuberculosis, including aggravated forms of TB such as DR-TB, MDR-TB and possibly XDR-TB. [23] Previous studies using Sanger sequencing protocols have shown that Mexico presents a considerable diversity of mutations associated with DR-TB, as well as variations related to geographical regions [24–31], which act to limit the utility of new genotypic diagnostic procedures and make WGS a very attractive alternative. This study therefore attempts to assess the diagnostic value of WGS for the detection of resistance to first line anti-tuberculosis drugs in isolates circulating in a region of Mexico with a high incidence of DR-TB.

## Materials and Methods

### Recovery of mycobacterial strains, culture and drug susceptibility test

An initial collection of 91 isolates of mycobacterial strains, with positive drug resistance determined by the phenotypic test method (BACTEC, MGIT 960 Becton-Dickinson) from 2012 to 2016, was randomly selected from the strain bank of the Public Health Institute of Veracruz. Demographic information of the patients bearing the selected strains was obtained from clinical laboratory records and anonymised.

*Mycobacterium tuberculosis* strains were isolated in Lowenstein-Jensen media, and sensitivity testing against first-line drugs was performed by the fluorometric method (BACTEC MGIT 960, Becton-Dickinson) using the following critical concentrations: isoniazid (H) 0.1μg/mL, rifampin (R) 1.0 μg/mL, ethambutol (E) 5.0 μg/mL and streptomycin (S) 1.0 μg/mL. Pyrazinamide (Z) sensitivity was determined using BACTEC MGIT 960 PZA kit (Becton Dickinson).

### DNA extraction, WGS sequencing and bioinformatics analysis

Extraction and purification of the genomic DNA from isolates was carried out following Van Soolingen et al. [32]. Quantification was conducted initially with a nanodrop (ThermoScientific, USA) with subsequent adjustment to a concentration of 0.2 ng/µL. The libraries were prepared according to the Nextera XT (Illumina, CA., USA) protocol, using 1ng of DNA quantified by Qubit fluorometer (Invitrogen, CA, USA). Quality was determined using TapeStation (Agilent Genomics) and the pool was normalized and loaded into a 300-cycle mid output cartridge at 1.8pM. Sequencing was performed using NexSeq 500 (Illumina, CA., USA), in a 2 × 150 paired-end format.

Due to the possible presence of DNA material that is not of the *M. tuberculosis* complex, Kraken software V2 [33] was used to classify the WGS reads and identify other Mycobacteria species. We further focused only on those reads that belong to the *Mycobacterium tuberculosis* species. The WGS analysis, including mapping and variant calling (SNP and INDELS), was performed following a previous pipeline [34]. Briefly, sequencing reads were mapped and aligned to an inferred MTBC most likely common ancestor genome using BWA. Variants were then separated into INDELS (small insertions and deletions included) and single nucleotide polymorphism (SNPs) using VarScan. In order to detect drug resistance-associated mutations with different frequencies within the population, single polymorphisms with at least 10 reads in both strands and a quality score of 20 were classified into two categories. We considered those mutations that involved a frequency range of 10 and 89% as variant-SNPs, and those with at least 90% frequency in the sample as constant-SNPs. An INDEL was considered where the mutation was present with at least 10x depth coverage. Variant annotation was performed using *M. tuberculosis* H37Rv annotation reference (AL123456.2). In addition, SNPs annotated in regions that were difficult to map, such as repetitive sequences and PPE/PE-PGRS genes, as well as those detected near INDELS, were excluded from the analysis.

### Identification of drug resistance-associated variants

In order to determine mutations related to drug resistance in the isolates included in the study, SNP data obtained by WGS were compared to a list of genes known to confer resistance to each anti-tuberculosis drug [35]: For resistance to rifampicin: *rpoB, rpoA* and *rpoC* genes; to streptomycin: *rrs, rpsl and gidB*; to isoniazid: *katG, inhA, oxyR-ahpC, ndh, fadE24 and FabG1*; to pyrazinamide: *pncA* and *rpsA*; to ethambutol: *embC, embA and embB*. For resistance to second-line drugs; Flouroquinolone: *gyrA* and *gyrB*; Aminoglycoside: *rrs* and *tlyA*; Ethionamide: *ethR, mshA, inhA, ndh ethA* and *fabG*; Linezolid: *Rrl* and *rplC*; Kanamycin/ethionamide: *eis*; p-amino salicylic acid: *thyA*; Isoniazid/ethionamide: *inhA;* and finally, for resistance to isoniazide, ethionamide and amikacin: *ndh*.

### Phylogenetic analysis and identification of transmission clusters

In order to classify the isolates, and to identify whether they had a specific genotype and whether these were related, a phylogenetic analysis was performed. We classified all of the strains according to the presence of the specific lineages or sub-lineages of phylogenetic variants proposed by Coll et al. 2014 [36] and Stucki et al. 2016 [37]. In order to detect highly likely genomic transmission clusters, we performed a pairwise genetic distance analysis based on a concatenated SNP alignment obtained from every constant-SNP from every isolate. A ≤12 SNP threshold was applied to delineate the transmission clusters, as proposed by Walker et al. 2013 [38]. A phylogeny including all the clinical isolates, with three samples used as an outgroup (one belonging to Lineage (L) 5, another to L6 and *Mycobacterium canetti* [34]), was inferred using the maximum likelihood phylogenetic approach implemented in MEGA V6 [39] by applying a general time reversible model of nucleotide substitution, with a gamma distribution (GTRGAMMA) and considering 1000 bootstraps. The tree was visualized in iTOL v. 4 (https://itol.embl.de/tree).

### Analysis of the diagnostic efficiency of WGS

Drug resistant profiles for each first-line drug, as predicted by WGS, were compared to the BACTEC MGIT phenotypic assay that was used as a reference standard for the diagnostic. The sensitivity (Sen), specificity (Spe), positive predictive values (PPV) and negative predictive values (NPV) were determined with 95% confidence intervals. Data was analyzed using STATA V.13. Inter-test agreement for both results was expressed by the percentage of overall agreement and kappa (κ) statistic, according to Landis et al. [40].

### Ethical concerns

No physical interventions took place with the patients and all of the information collected was treated as confidential. The ethics committee of the Public Health Institute at University of Veracruz oversaw and approved the ethical issues involved in this study

## Results

### Isolate recovery, patient data and resistotyping

Ninety-one clinical isolates, obtained from the same number of individuals with TB and resistance to at least one first-line drug, were initially included in the study. Further genome analysis confirmed the presence of the species *M. tuberculosis* in 81 of the isolates. Four isolates presented *M. fortuitum* and two presented *M. abscessus* and it was not possible to determine the species in a further four isolates. These ten isolates were therefore excluded from the global study.

Of the 81 individuals ultimately included in the study, 50% (42) were male, with a mean age of 45 years (±14.6) and all individuals presented respiratory TB. A total of 36% (31) presented type 2 diabetes mellitus (T2DM), 47% (40) were undergoing primary treatment and 42% (36) presented a re-treatment.

In relation to the phenotypic drug resistance profile of the 81 isolates determined by BACTEC MGIT assay, S resistance was observed in 47% (38) of the isolates, while H resistance was presented in 75% (61), R resistance in 62% (n=51), E resistance in 29% (24) and Z resistance in 34% (28). A total of 25% (21) of the isolates were classified as mono-resistant, 14% (12) as poly-resistant, 56% (46) as MDR-TB and 16% (13) as resistant to all first-line drugs.

### Polymorphism conferring resistance against first-line drugs

According to the BACTEC MGIT phenotypic assay, 61 isolates showed H resistance, of which 51 (62%) presented polymorphisms that could explain their condition, while no mutations were detected in ten (12%) of the isolates, according to the WGS analysis. The most frequent mutation identified was *katG315S/T* in 35 (60%) isolates, followed by the intergenic position 1673425 (c/t) of the *fabG1* gene in nine (14%) isolates and the intergenic positions 2726105088-2726145 of *ahpC-OxyR*, which were observed in seven (11%) isolates (Table 1).

**Table 1.** Isoniazid mutations obtained by WGS from *Mycobacterium tuberculosis* isolates circulating in Veracruz, Mexico

| Position | Region | Codon | Amino Acid | Change | Frequency |
| --- | --- | --- | --- | --- | --- |
| ***katG*** |  |  |  |  |  |
| 2154250<sup>1</sup> | Coding | 621 | A → V | gcc/gtc | 1* |
| 2155139<sup>1</sup> | Coding | 325 | P → S | ccg/tcg | 1 |
| 2155167 | Coding | 315 | S → R | agc/aga | 1 |
| 2155168 | Coding | 315 | S → T | agc/acc | 35 |
| 2155168 | Coding | 315 | S → I | agc/atc | 1* |
| 2155168 | Coding | 315 | S → N | agc/aac | 1 |
| 2155700 | Coding | 138 | N → H | aac/cac | 2* |
| ***ahpC-OxyR*** |  |  |  |  |  |
| 2726105 | Intergenic | --- | --- | g/a | 2* |
| 2726117 | Intergenic | --- | --- | t/a | 2* |
| 2726139 | Intergenic | --- | --- | c/t | 1 |
| 2726145 | Intergenic | --- | --- | g/a | 2 |
| ***fabG1*** |  |  |  |  |  |
| 1673425 | Intergenic | --- | --- | c/t | 9 |
| Total |  |  |  |  | 58* |

\* In combination with other mutation in a single isolate.

Of the 51 isolates that were rifampicin-resistant according to the BACTEC MGIT phenotypic assay, 49 (96%) had a mutation in the *rpoB* gene. Some isolates presented more than one mutation, while two isolates presented no mutations. One sensitive isolate, as determined by the BACTEC assay, showed the *rpoB430L/P* mutation while another strain had the mutation *rpoB432Q/L*. Thirteen mutations were observed, the most frequent of which was found at *rpoB450S/L* in 35 (71%) of the isolates (Table 2).

**Table 2.**
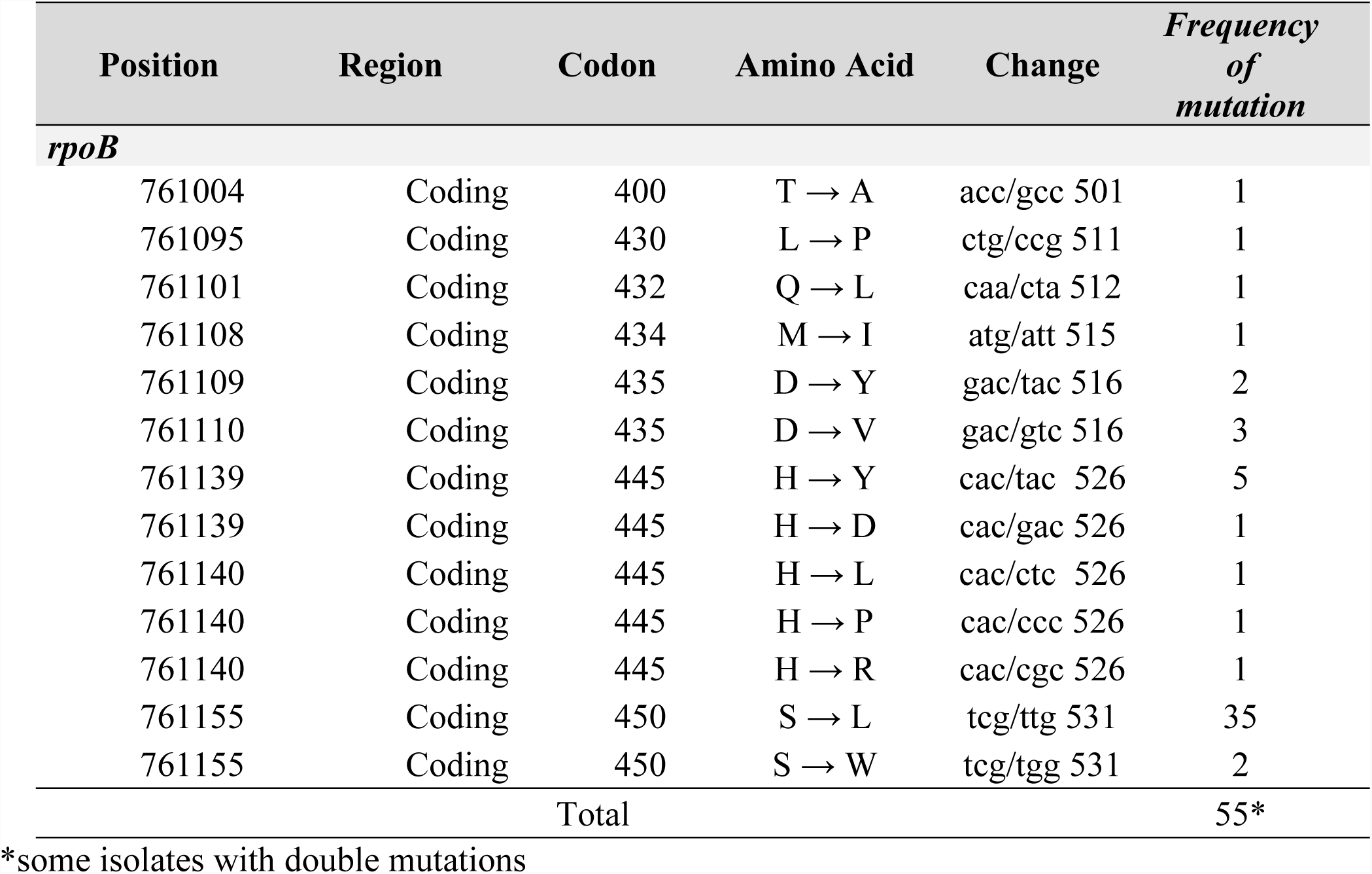
Rifampicin mutations detected by WGS from isolates of *Mycobacterium tuberculosis* circulating in Veracruz, Mexico

Of the 24 isolates with phenotypically E resistance, 17 (71%) presented polymorphisms that explained their condition, according to WGS. However, seven (30%) of the resistant isolates showed no mutation, mine while six isolates, defined as sensitive by the BACTEC MGIT phenotypic assay, showed one mutation related to resistance to this drug. The most frequent mutation was observed at *embB306M/V/I* in eleven isolates, followed by the *embA* promoter C-12T mutation at the intergenic genome position 4243221, which was observed in eight isolates (Table 3).

**Table 3.**
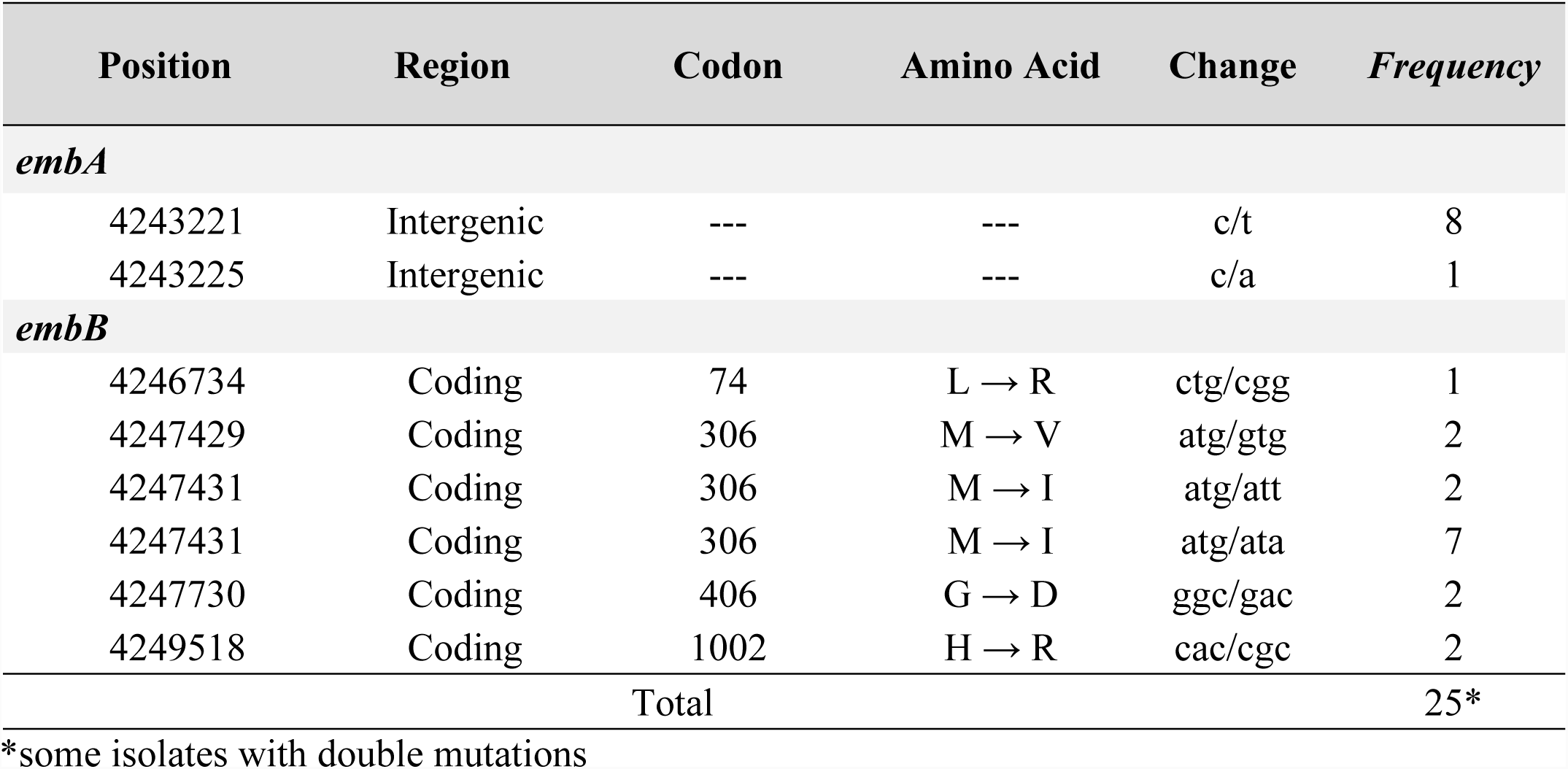
Ethambutol mutations obtained by WGS from isolates of *Mycobacterium tuberculosis* circulating in Veracruz, Mexico

| Position | Region | Codon | Amino Acid | Change | Frequency |
| --- | --- | --- | --- | --- | --- |
| ***embA*** |  |  |  |  |  |
| 4243221 | Intergenic | --- | --- | c/t | 8 |
| 4243225 | Intergenic | --- | --- | c/a | 1 |
| ***embB*** |  |  |  |  |  |
| 4246734 | Coding | 74 | L → R | ctg/cgg | 1 |
| 4247429 | Coding | 306 | M → V | atg/gtg | 2 |
| 4247431 | Coding | 306 | M → I | atg/att | 2 |
| 4247431 | Coding | 306 | M → I | atg/ata | 7 |
| 4247730 | Coding | 406 | G → D | ggc/gac | 2 |
| 4249518 | Coding | 1002 | H → R | cac/cgc | 2 |
| Total |  |  |  |  | 25* |

\*some isolates with double mutations

Of the 28 isolates with resistance to Z according to the phenotypic assay, 21 (75%) had mutations at *pncA* while seven presented a wild-type genotype. Four isolates phenotypically identified as Z-susceptible presented a mutation that could be associated with resistance. In total, 17 mutations were identified in *pncA*. The most frequent of these was *pncA120L/P*, which was detected in 14 isolates (Table 4).

**Table 4.**
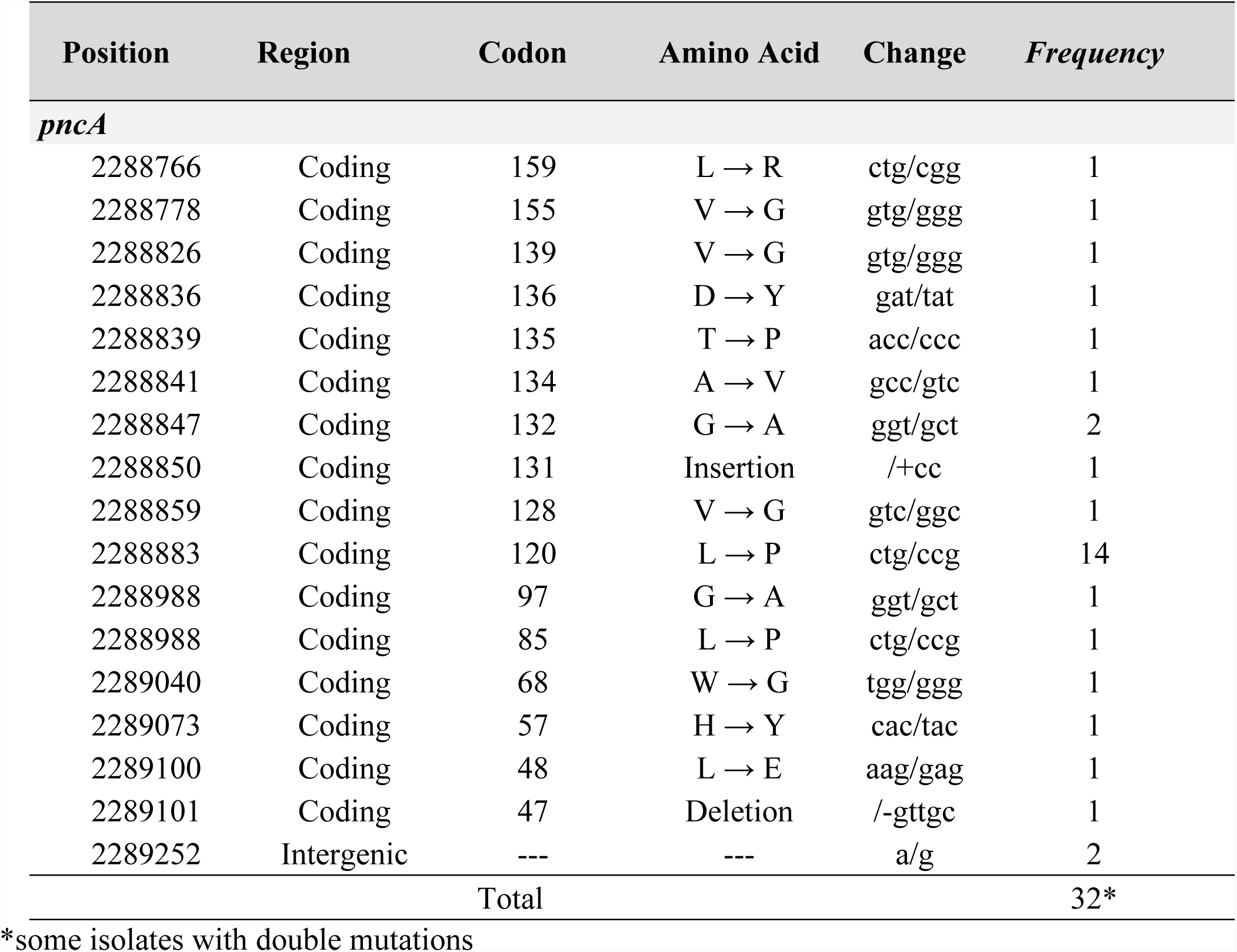
Pyrazinamide mutations obtained by WGS from isolates of *Mycobacterium tuberculosis* circulating in Veracruz, Mexico

Of the 38 isolates with phenotypic S resistance, only eleven (29%) presented a mutation that would explain the resistant condition (Table 5). The remaining 27 (71%) isolates presented no change. Only one phenotypically-determined sensitive isolate showed one polymorphism associated with resistance. Nine mutations were observed in three genes (*gidB, rpsL* and *rrs*); the most abundant of which were *rpsL43K/R* and a c/t mutation at position 1472362 on the *rrs* gene, both found in three isolates each. (Table 5)

**Table 5.** Streptomycin mutations obtained by WGS from isolates of *Mycobacterium tuberculosis* circulating in Veracruz, Mexico

| Position | Region | Codon | Amino Acid | Change | Frequency |
| --- | --- | --- | --- | --- | --- |
| ***gidB*** |  |  |  |  |  |
| 4407790 | Coding | 138 | A → V | gcg/gtg | 1 |
| 4407992 | Coding | 71 | G → R | gga/aga | 1 |
| ***rpsL*** |  |  |  |  |  |
| 781687 | Coding | 43 | K → R | aag/agg | 3 |
| 781821 | Coding | 88 | K → Q | aag/cag | 1 |
| 781822 | Coding | 88 | K → T | aag/acg | 1 |
| 781822 | Coding | 88 | K → R | aag/agg | 1 |
| ***rrs*** |  |  |  |  |  |
| 1472359 | Ribosomal | --- | --- | a/c | 1 |
| 1472362 | Ribosomal | --- | --- | c/t | 3 |
| 1472751 | Ribosomal | --- | --- | a/g | 1 |
| Total |  |  |  |  | 13* |

\*some isolates with double mutations

### Polymorphism conferring multidrug and second line drug resistance

Of the 46 isolates identified as MDR-TB by the BACTEC MGIT method, 41 (89%) showed simultaneous mutations in the genes associated with resistance against rifampicin and isoniazid by WGS. Four isolates (7%) lacked any polymorphism associated with isoniazid resistance in the *katG, inhA, oxyR-ahpC* or *FabG1* genes analyzed, and two isolates (3%) did not present a mutation associated with rifampicin resistance in *rpoB, rpoA* and *rpoC* genes.

It was not possible to determine phenotypic resistance against second-line drugs in the isolates analyzed because of limitations in reagents and infrastructure. However, the WGS analysis of genes related with resistance against these drugs showed that twelve isolates presented a mutation that would confer resistance to fluoroquinolones; ten had a mutation at *gyrA*, conferring resistance to ofloxacin, and two had one mutation at *gyrB*, conferring resistance to levofloxacin (Table 6). In addition, five isolates presented mutations at *eis* and *rrs*, conferring resistance to kanamycin, amikacin and capreomycin. Finally, no polymorphisms were observed in genes associated with resistance to linezolid (*rrl* and *rplC*), aminoglycosides (*tlyA*), ethionamide (*ethR, mshA, inhA, ndh* and *ethA*) and p-amino salicylic acid (*thyA*).

**Table 6.** Second line drug mutations determined through WGS from isolates of *Mycobacterium tuberculosis* circulating in Veracruz, Mexico.

| Position | Region | Codon | Amino Acid | Change | Frequency |
| --- | --- | --- | --- | --- | --- |
| **Fluoroquinolones** |  |  |  |  |  |
| ***gyrB*** |  |  |  |  |  |
| 6738 | Coding | 500 | T → N | acc/aac | 2 |
| ***gyrA*** |  |  |  |  |  |
| 7564 | Coding | 88 | G → A | ggc/gcc | 1 |
| 7570 | Coding | 90 | A → V | gcg/gtg | 3 |
| 7581 | Coding | 94 | D → Y | gac/tac | 1 |
| 7582 | Coding | 94 | D → A | gac/gcc | 3 |
| 7582 | Coding | 94 | D → G | gac/ggc | 2 |
| Total |  |  |  |  | 12 |
| **Kanamycin (KM) / Amikacin (AM) / Capreomicin (CM)** |  |  |  |  |  |
| ***Eis*** |  |  |  |  |  |
| 2715342 | Intergenic | --- | --- | g/a | 1 |
| ***Rrs*** |  |  |  |  |  |
| 1473246 | Ribosomal | --- | --- | a/g | 4 |
| Total |  |  |  |  | 10 |

\*some isolates with double mutations

A joint analysis of MDR-TB isolates with the mutations related to resistance against second-line drugs showed the presence of eight isolates that could potentially be considered as pre-XDR-TB (with resistance to H and R, associated with resistance to FQ or a second-line injectable, but not both), as well as three isolates that were recognized as XDR-TB (with resistance to H and R, and resistance to any of the fluoroquinolones (such as levofloxacin or moxifloxacin)), and to at least one of the three injectable second-line drugs (amikacin, capreomycin or kanamycin).

### Concordance between WGS mutations and phenotypic resistance and kappa test

Considering the diagnosis obtained by BACTEC MGIT phenotypic assay as the reference test, the efficacy of WGS in terms of predicting resistance against H, R, E, Z and S showed sensitivity values of 84%, 96%, 71%, 75% and 29% and specificity values of 100%, 94%, 90%, 90% and 98%, respectively. The positive predictive values (PPV) were 100%, 96%, 74%, 84% and 92%, while the negative predictive values (NPV) were 67%, 94%, 89%, 85% and 62%, respectively. Finally, for multi-drug resistance, sensitivity and specificity values of 89% and 97%, and PPV and NPV values of 97% and 88%, respectively, were determined.

The analysis of concordance with Cohen’s kappa test between the phenotypic assay and the resistance profile determined by WGS showed substantial agreement for H (k: 0.73, 88%), perfect agreement for R(k=0.90, 95%), substantial agreement for E (k=0.61, 84%) and Z (k= 0.67, 84%), fair agreement for S (k= 0.28, 66%) and almost perfect agreement for MDR-TB (k= 0.85, 92%).

### Phylogenetic analysis and identification of transmission cluster

Four major lineages (L) were observed through the WGS analysis. Two (2.3%) isolates were included in L1 (EAI), both of which were MDR-TB. Four isolates (5%) were located within L2 (Beijing) and included three MDR-TB isolates and one potential pre-XDR-TB isolate. Two isolates (2.3%) were located inside the L3 (CAS), both considered through the WGS as potentially pre-XDR-TB. The remaining 76 (90%) isolates were classified within L4, and included eight sub-lineages (T, T1, X2, X3-X1, LAM3, LAM9, LAM11 and S). The most abundant was sub-lineage 4.1.1 (X), which was present in 23 (28%) isolates, followed by sub-lineage 4.1.2 (H) found in 22 (27%) isolates and sub-lineage 4.3.3 (LAM) found in eight (10%) isolates (figure 1).

**Figure 1.**
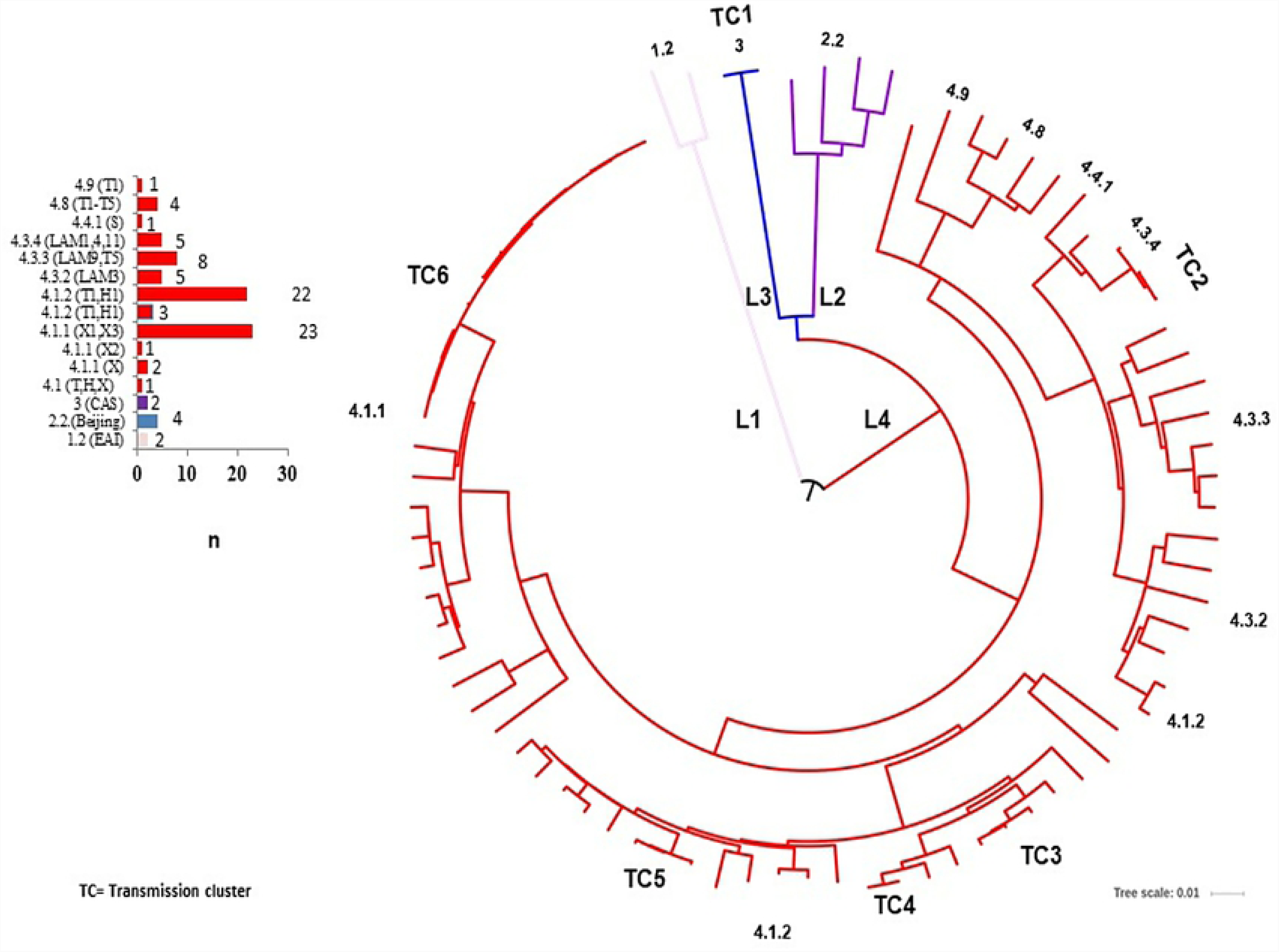
Phylogeny and transmission clusters of *M. tuberculosis* isolates from Veracruz, Mexico obtained by WGS

The WGS analysis allowed identification of six transmission clusters (TC); TC1 included two isolates with L3, also sharing the polymorphisms related to a pre-XDR-TB character. The rest of the TC were located within L4; TC2 included three isolates with sub-lineage 4.3 (LAM), TC3 included three isolates, TC4 had two isolates and TC5 had three isolates, all included in the sub-lineage 4.1.2 (H). Finally, TC6 included thirteen isolates with sub-lineage 4.1.1 (X1-X3), all sharing a multidrug resistant condition. One presented the polymorphisms that could potentially classify it as pre-XDR-TB and another as XDR-TB (Figure 1).

## Discussion

Mexico is a country that presents endemic tuberculosis and more than 20,000 cases are reported annually, of which close to 1,500 are DR-TB. The state of Veracruz has a population of close to seven million inhabitants, and contributes 13% (2,500) of the total number of TB cases in Mexico, close to 15% of the annually reported number of DR-TB cases in the country and occupies first place in terms of prevalent cases of MDR-TB [22]. For these reasons, Veracruz is positioned as one of the most significant contributors to the DR-TB and MDR-TB problem in Mexico. The epidemiological characteristics of the population included in this study, such as a high proportion of individuals with T2DM (36%) and primary treatment (47%), reflect previous reports for the region [24,25,30,41] and confirm the influence of T2DM as an important factor in the development of DR-TB and MDR-TB, as well as highlighting the difficulties for diagnosis, control and preventive management of DR-TB by the health agencies as well as evidencing the urgent need to develop new diagnostic procedures.

Among the 81 DR-TB isolates subjected to WGS, more than 200 polymorphisms were identified in 60 positions of the more than 35 genes analyzed. This evidences the diversity of mutations present in the drug-resistant isolates in circulation, and is in agreement with previous reports describing mutations identified by Sanger-type sequencing [27–30,42–44].

A sensitivity of 84%, specificity of 100% and kappa concordance of 88% were determined for the diagnosis of resistance to H; these values are similar to that described in reports from different locations [8–11,21] (Table. 7). Ten isolates showed no mutations that could explain their phenotypic condition of resistance. The possibility exists that other mutations or unreported genes, such as pump flux, may be actively participating but further studies are required to confirm this possibility. There was a noteworthy presence of isolated carriers of unusual mutations, such as *katG*P325S and *katG*N138H [45], as well as an important number of mutations in the intergenic position of ahpC-OxyR, described as compensatory for isoniazid resistance and associated with *katG* mutations [46]. Considering all the above, the information obtained supports the proposal of the use of WGS as a useful tool for the diagnosis of isoniazid resistance [46].

**Table 7.**
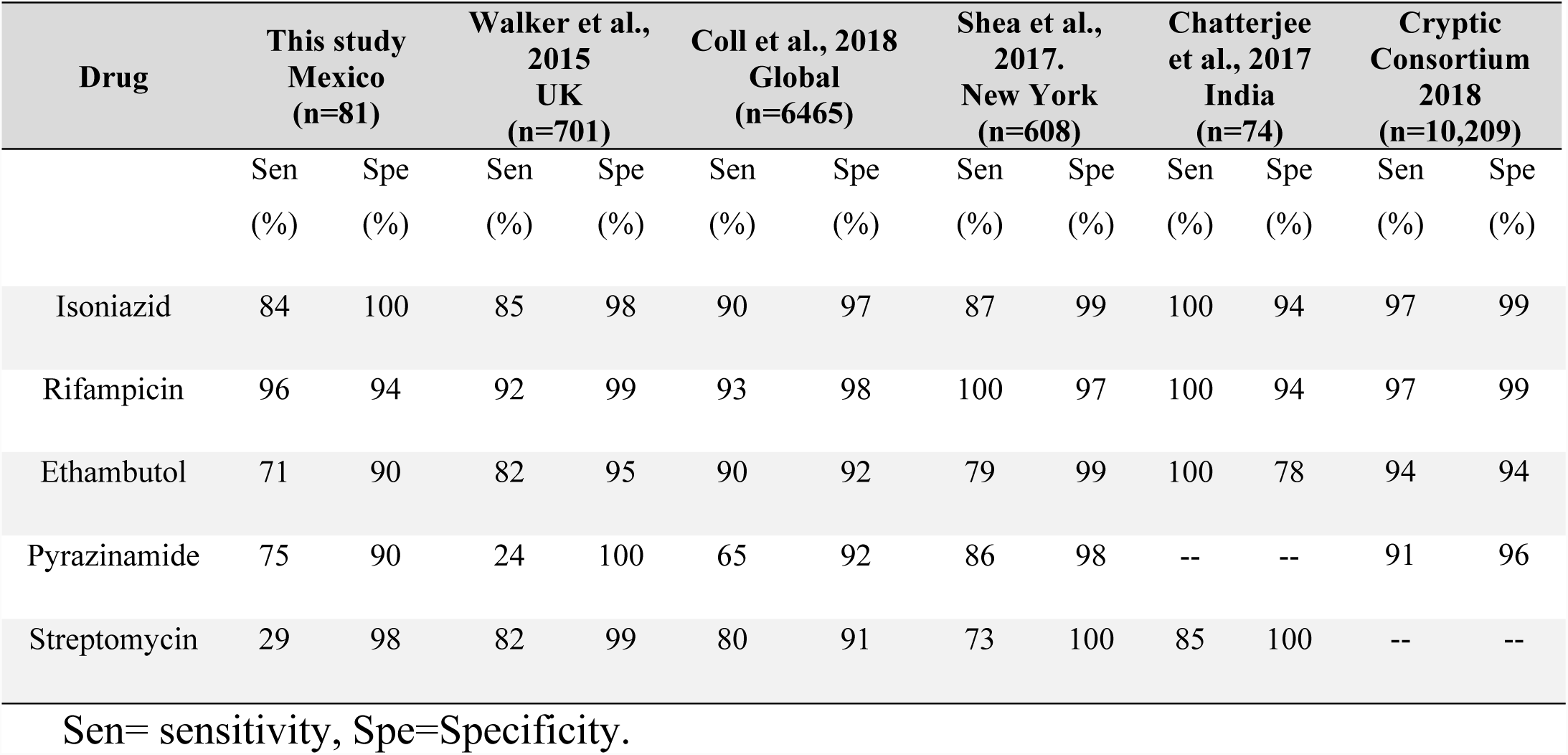
Comparison of percentage of concordance values for drug resistant diagnosis prediction obtained by WGS found in this and several other studies

| Drug | This study Mexico (n=81) |  | Walker et al., 2015 UK (n=701) |  | Coll et al., 2018 Global (n=6465) |  | Shea et al., 2017. New York (n=608) |  | Chatterjee et al., 2017 India (n=74) |  | Cryptic Consortium 2018 (n=10,209) |  |
| --- | --- | --- | --- | --- | --- | --- | --- | --- | --- | --- | --- | --- |
|  | Sen | Spe | Sen | Spe | Sen | Spe | Sen | Spe | Sen | Spe | Sen | Spe |
|  | (%) | (%) | (%) | (%) | (%) | (%) | (%) | (%) | (%) | (%) | (%) | (%) |
| Isoniazid | 84 | 100 | 85 | 98 | 90 | 97 | 87 | 99 | 100 | 94 | 97 | 99 |
| Rifampicin | 96 | 94 | 92 | 99 | 93 | 98 | 100 | 97 | 100 | 94 | 97 | 99 |
| Ethambutol | 71 | 90 | 82 | 95 | 90 | 92 | 79 | 99 | 100 | 78 | 94 | 94 |
| Pyrazinamide | 75 | 90 | 24 | 100 | 65 | 92 | 86 | 98 | -- | -- | 91 | 96 |
| Streptomycin | 29 | 98 | 82 | 99 | 80 | 91 | 73 | 100 | 85 | 100 | -- | -- |

Sen= sensitivity, Spe=Specificity.

The sensitivity of 96% and specificity of 94% observed for the diagnostic of R by WGS coincides with previous reports [8–11,21] (Table 7). Two isolates were notable for being phenotypically susceptible to rifampicin, despite the fact that they were carriers of the mutations *rpoB432Q*/L and *rpoB430L/P* respectively, which have been considered as “disputed mutations” [47] that confers low-level R resistance [48]. Isolates carrying this type of mutation are phenotypically misclassified as rifampicin susceptible, and routine genotypic tests fail in terms of their detection [47]. This indicates that WGS could improve the local diagnostic of R resistance, and of MDR, considering that rifampicin resistance has become a subrogated marker of MDR-TB [1]. In this context, the analyses of the 51 R-resistant isolates included in this study show that 46 (90%) of these were also MDR-TB according to the BACTEC MGIT phenotypic test, while 41 (80%) presented, through WGS, the mutations that explain that condition.

The sensitivity for the diagnosis of resistance to E by WGS was 71%, which is one of the lowest described to date [8–11,21] (Table 7). Seven resistant isolates showed no mutations, while six sensitive isolates showed a mutation associated with resistance to this drug. These contradictory results can explain the low sensitivity and positive predictive values observed. Such variations in the occurrence of mutations associated with resistance to E are similar to those reported previously [49]. Resistant isolates lacking in mutations and the presence of mutations related to resistance in sensitive isolates show the possibility of failures in the BACTEC MGIT phenotypic assay, and further analysis is required in order to identify the factors that cause this low sensitivity and to improve the efficiency of WGS in predicting resistance against this drug among the isolates from the setting.

The sensitivity of 75% observed for Z was low compared to other reports [8–11,21] (Table 7); however, the specificity of 90% and kappa agreement of 84% supports the use of WGS for the diagnosis of resistance to this drug. The presence of four susceptible but mutation-carrying isolates and seven resistant but unmutated (*pncA or rpsA*) isolates can help to explain the low sensitivity found. However, technical problems with *in vitro* testing of Z resistance associated with inoculum concentration, volume and homogeneity [50], and even changes in the pH of the culture medium, have been considered to influence the diagnostic of false positive resistant Z strains [51]. The diversity and number of mutations and indels observed in *pncA* was the highest among all of the genes studied, and was similar to values previously described [29]. A deletion in the *pncA47* position (2289101; - gttgc) and an insertion in *pncA131* (2288850; +cc) were identified; both indels were described in isolates circulating in the region five years ago [29], raising the possibility that these isolates are still actively circulating in the present population.

The 29% sensitivity observed for S through the WGS was among the lowest reported [8–11,21] (Table 7). The mechanisms of resistance to S are complex and involve several genes. While those considered as canonical (*gibB, rpsL and rrs*) were characterized here, only a limited number of isolates carried a mutation; a behavior similar to that previously observed in isolates from the region [28], highlighting the need for further studies to identify the mutations involved with resistance against this drug.

While S has not been used in Mexico as a first line drug against TB since the 1980s, the fact that 47% of the isolates analyzed presented resistance to this drug evidences the possible presence of “old” S resistant strains still in active circulation in the population and that actually participate in the resistance against first line drugs in TB. This could be of particular relevance in Mexico, considering the WHO recommendation for use of this drug as a second line drug in specific situations [52].

The sensitivity and specificity for the diagnosis of MDR-TB using WGS were 89% and 97%, respectively, and presented a kappa agreement of 92%. These values indicate the feasibility of using this technique as a support for prediction of the diagnosis of these aggravated forms of TB. There is no doubt that this will have a significant value in Mexico, since it is a country that generates an important number of cases of MDR-TB in Latin America.

Through WGS, eight isolates were identified as bearing the mutations that would confer a potential pre-XDR-TB character, and three isolates were predicted as XDR-TB; nevertheless, it was not possible to acquire their phenotypic profiles against second line drugs and confirm the real condition of these isolates. In this sense, previous reports describe that it is possible by means of WGS to identify XDR-TB isolates with sensitivity and specificity values exceeding 85% [6,8,10,18,53]. Thus, the possibility of generating a faster screening diagnosis of pre- and XDR-TB strains using WGS could have important implications for a middle-income country such as Mexico, where at least 6 to 12 months are required to establish a sensitivity profile against second line drugs in isolates suspected of having this condition.

While it was only a relatively small number of isolates studied, four lineages were identified; 90% (76) were placed in L4 (Euro-American), confirming the predominance of this lineage in the population [24,25,41,54,55] and the presence of the “founder effect”, where the predominance of specific lineages seems to limit the dispersion of new or less frequent lineages [55–57]. Twenty-six (32%) isolates were located within six transmission clusters, of which two were particularly noteworthy: i) the TC1, which included the only two isolates with L3 (CAS) and ii) the TC6, comprising 13 (17%) MDR-TB isolates with sub-lineage 4.1.1 (X), including one isolate identified as pre-XDR-TB and another as XDR-TB. These CAS and X lineages are rarely referred in Mexico, particularly with such a strong tendency towards drug resistance [24,25,41,54,58] and additional studies will be necessary to determine the extension of this sub-lineages in the population.

The main limitation of this study probably relates to the fact that all of the isolates came from a single region of Mexico, even though it is one of the states with the highest number of cases of DR-TB and MDR-TB in the country. It would undoubtedly be necessary to increase the number of isolates sourced from different regions of the country, and to identify the mutations that are most frequently involved in the development of resistance to first- and second-line drugs. This information will guide the application of WGS, for early diagnosis and treatment of DR-TB, as well as the development of epidemiological studies to identify the risk factors associated with the development and transmission of infection generated by DR-TB in the Mexican population.

In conclusion, this study illustrates the utility of WGS in terms of the diagnostic of resistance against first line drugs, MDR-TB and the screening of pre-XDR- and XDR-TB in Mexico; as well as presenting the valuable additional possibility of characterizing the genotypes in circulation and identifying transmission clusters, allowing further development of epidemiological-genomic surveillance studies.

## Acknowledgments

CFM-M was a CONACYT fellow of the Maestría en Ciencias de la Salud of the Instituto de Ciencias de la Salud Universidad Veracruzana. RZ-C and IC were partially funded by I-COOP-2017-COOPB2032. RMS was funded by CONACyT-Problemas nacionales grant no. 2015-01-147. XS was funded by GACD-FONCICYT Conacyt No.264693. Thanks go to Nancy Seraphín, from the Division of Infectious Diseases and Global Medicine, University of Florida, Gainesville, Florida, USA, for performing the WGS of certain isolates.

